# Dopamine and calcium dynamics in the nucleus accumbens core during food seeking

**DOI:** 10.1101/2025.03.11.642710

**Authors:** Sophia J. Weber, Gillian S. Driscoll, Madelyn M. Beutler, Hayley M. Kuhn, Jonathan G. Westlake, Marina E. Wolf

## Abstract

Extinction-reinstatement paradigms have been used to study reward seeking for both food and drug rewards. The nucleus accumbens is of particular interest in reinstatement due to its ability to energize motivated behavior. Indeed, previous work has demonstrated that suppression of neuronal activity or dopaminergic signaling in the nucleus accumbens reduces reinstatement to food seeking. In this study, we sought to further establish a connection between glutamatergic input, measured by proxy via a genetically encoded calcium indicator, and dopamine (DA) tone, measured simultaneously with a red-shifted DA biosensor. We performed this sensor multiplexing in the nucleus accumbens core in the classic extinction-reinstatement paradigm with food reward. We detected DA transients that changed in magnitude and/or temporally shifted over the course of self-administration training. In our calcium traces we observed a decrease from baseline time-locked to the lever press for food reward, which became more prominent with training. Both patterns were reduced in the first session of extinction with no deflections from baseline detected in either the DA or calcium traces in the last extinction session. When we recorded during reinstatement tests, bootstrapping analysis detected a calcium response when reinstatement was primed by cue or pellet+cue presentation, while a DA response was detected for pellet+cue reinstatement. These data further establish a role for nucleus accumbens core activity and DA in reinstatement of food seeking and represent the first attempt to simultaneously record the two during an extinction-reinstatement task.

## Introduction

Reward-associated stimuli help guide motivated behavior, and these learned associations between an environmental cue and rewarded outcome are generally adaptive. However, the ability of reward-associated cues to elicit behavior can become problematic when these rewards also result in negative consequences, as with addictive drugs. Reinstatement of reward seeking has been studied for many decades to interrogate drivers of maladaptive reward seeking behavior, particularly as an analog for relapse in substance use disorder (Shaham et al., 2003; Venniro et al., 2016). However, this model has also been used with palatable food reward instead of drug reward to study mechanisms of food seeking (Calu et al., 2014; Scofield et al., 2016). It allows for the examination of neurobiological circuits of reward learning across multiple phases: acquisition, extinction, and reinstatement.

The nucleus accumbens (NAc) is an important locus for goal-directed behavior, integrating information from cortical, limbic, and midbrain inputs and sending output to motor regions (Floresco, 2015). It has long been recognized as a particularly important region in food seeking and consummatory behavior (Kelley, 2004; Alonso-Caraballo et al., 2021). Early studies found that a subset of neurons in the NAc exhibit a pause in firing during initiation and maintenance of sucrose consumption (Nicola et al., 2004; Roitman et al., 2005; Taha and Fields, 2005, 2006), with a more study showing that electrical stimulation of NAc regions containing such neurons interrupts consumption (Krause et al., 2010). In operant tasks, recordings of NAc neurons also reveal subpopulations of MSN with distinct response patterns. For example, a subset of phasically active MSN exhibits an increase in firing prior to lever pressing for food or water reward with a separate subset showing inhibition immediately before and after responding (Carelli et al., 2000). Furthermore, activity of NAc MSN regulates food seeking in extinction-reinstatement paradigms. For example, pharmacological inactivation of the NAc core (NAcc), but not shell, has been shown to impair cue-induced reinstatement (Floresco et al., 2008).

At the level of neurotransmitters in the NAc, dopamine (DA) transmission is important for reward seeking in food tasks (Volkow et al., 2017; Marinescu and Labouesse, 2024) despite not being necessary for the hedonic value of reward (Berridge and Robinson, 1998). For example, blockade of either DA D1 receptors or DA D2 receptors in the NAcc inhibited food reinstatement (Guy et al., 2011). More recently, a series of studies have established the importance of plasticity of glutamate transmission in the NAc for regulation of food seeking (Ferrario, 2020). These findings suggest that DA and glutamatergic input into the NAcc may converge to facilitate food reinstatement, which is supported by evidence for these systems working together in other circumstances (Nicola et al., 2000; Wolf, 2010; Tritsch and Sabatini, 2012; Sippy and Tritsch, 2023; Weber et al., 2024a).

We sought to establish a fiber photometry approach capable of more directly interrogating the relationship between glutamate and DA signaling during a food extinction-reinstatement paradigm. To do so, we employed fiber photometry with sensor multiplexing to simultaneously record NAcc DA release and MSN calcium levels, a proxy for glutamatergic activation, during all phases of a food extinction-reinstatement task. We found that DA release increased during the learning or self-administration phase, and that eventually this peak shifted earlier in time to precede the lever press. This DA signal then declined with extinction and re-emerged with reinstatement (based on bootstrapping analysis). In contrast, MSN calcium levels decreased from baseline after the lever press during self-administration. This decrease disappeared after the first day of extinction. Reinstatement was associated with an increase in MSN calcium and DA. These results are the first to examine both MSN activity via calcium proxy and DA release during an extinction-reinstatement task. Furthermore, this study paves the way for future multiplexing studies of addictive drugs, perhaps helping us better understand the relationship between DA release and postsynaptic MSN activity in addiction models.

## Methods

### Subjects

We used male and female Long-Evans rats obtained from Charles River (Wilmington, MA). They were ∼9 weeks old upon arrival and were group housed for ∼1 week after arrival to acclimate prior to surgery. After surgery, all rats were single housed. During recovery, rats had free access to standard laboratory chow and water in their home cages and were maintained on a reverse 12:12h light/dark cycle. One week prior to starting behavioral training, rats began food restriction and were maintained at 85% of their free-feeding weight, although water was freely accessible in the home cage throughout the experiment. Weights were recorded throughout food restriction. All procedures were approved by the OHSU Institutional Animal Care and Use Committee and followed NIH guidelines outlined in the Guide for the Care and Use of Laboratory Animals. From a total of 8 male and 7 female rats assigned to experimental groups, 4 were excluded after histology due to misplaced intracranial cannulas.

### Surgery

*Intracranial virus infusion and fiber optic cannula implantation:* We performed bilateral (n=8, 4 male/4 female; for these rats, one hemisphere was selected for recording) or unilateral (n=3, 2 male/1 female) intracranial infusions of viral cocktail of GCaMP8s, a genetically encoded calcium indicator (AAV9-syn-jGCaMP8s-WPRE, Addgene #162374), and the red-shifted dopamine biosensor rGRAB_DA3m (AAV2/9-hsyn-rDA3m, Biohippo #BHV12400545), into the NAcc followed by implantation of fiber optic cannula (Thor Labs, CFM15L10) above each infusion site. We performed unilateral surgeries on a subset of rats due to lack of supplies after shipping was delayed due to an ice storm. Briefly, rats were mounted in a stereotaxic device and, after making an incision, the nose bar was adjusted such that the change in dorsal-ventral coordinates from Lambda to Bregma was <0.1 mm. We performed two infusions per hemisphere (250 nL each at 100nL/min of working stock consisting of 3 µL GCaMP8s, 3 µL rGRAB_DA3m, and 3 µL sterile saline) using the following stereotaxic coordinates (relative to Bregma): AP +1.3; ML ±2.4 mm; DV-7.2 mm for first infusion and DV-7.0 mm for the second infusion, 6° angle (Paxinos and Watson, 2014). Following the virus infusion, we implanted fiber optic cannula (AP +1.3 mm; ML ±2.4 mm; DV-6.8 mm; 6° angle). We anchored the cannula to the skull with 1/8” pan head sheet metal screws (Fastenere #842176107226) and dental cement (Stoelting #51458) and covered the fiber optic cannula with dust caps (Thor Labs, CAPF) for protection. Rats received 5 mg/kg subcutaneous meloxicam (Covetrus, 6451602845, SKU #49755) as a post-operative analgesic. Rats then recovered for 7 days prior to beginning food restriction (see Fig. 1A for timeline).

**Figure 1.**
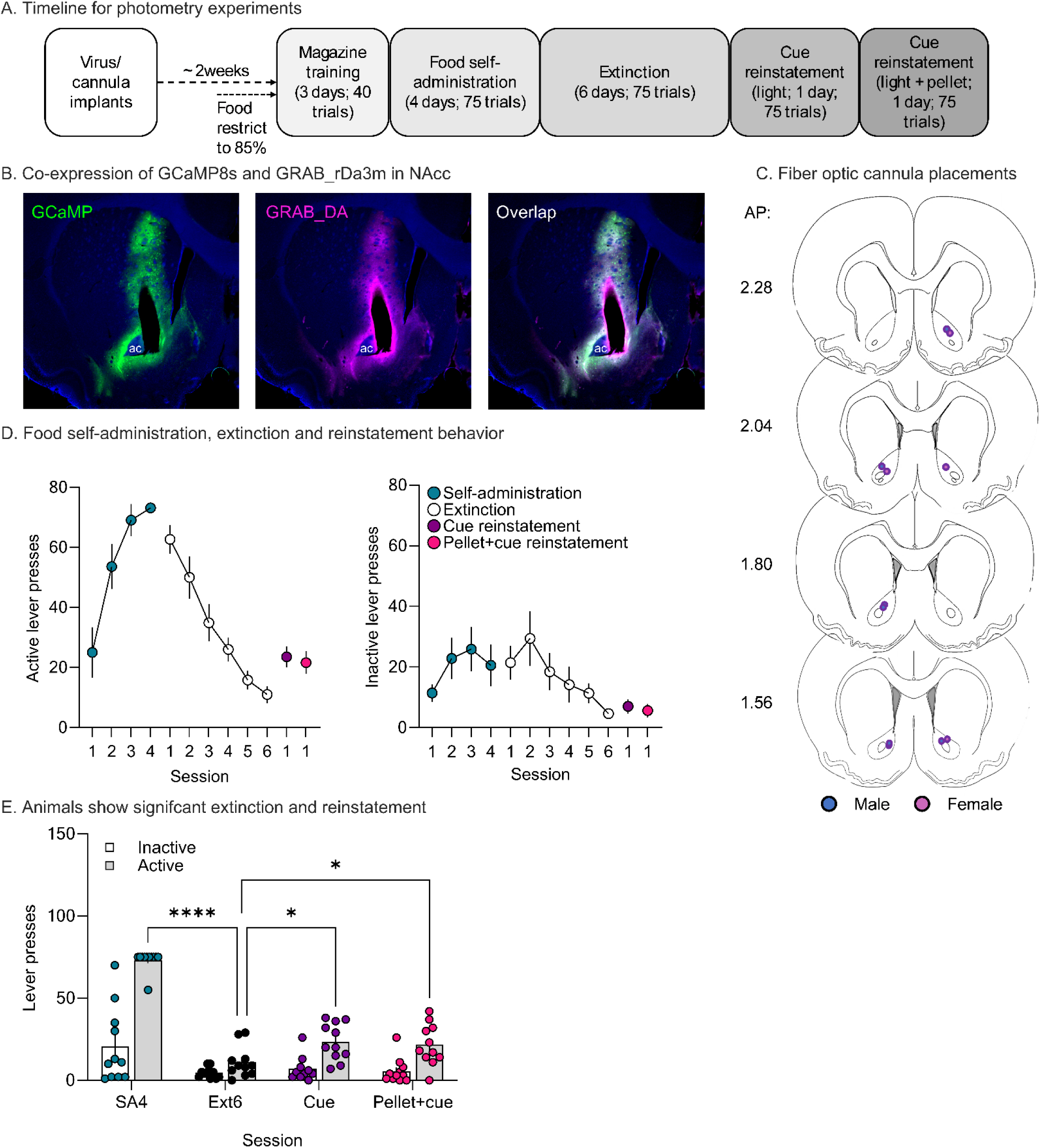
Behavior and virus expression. **A**. Timeline for experiment. **B**. Representative image of virus co-expression: *left,* AAV9-syn-jGCaMP8s-WPRE0, *middle,* AAV2/9-hsyn-rDA3m, and *right,* Overlap of viral expression. The cannula track appears as an oblong shape terminating to the right of the anterior commissure. **C**. Fiber optic cannula placement for rats. The dot signifies the end of the cannula (females, pink; males, blue). Placement was determined after immunohistochemistry for GFP and mApple to confirm virus expression at the end of the cannula. Sections adapted from Paxinos & Watson 7^th^ edition. **D**. Food self-administration sessions (SA1-4), extinction sessions (Ext1-6), and reinstatement tests. *Left:* Active lever presses for all rats (n = 11) across all sessions for all behavioral phases. *Right:* Inactive lever presses for the same sessions. **E.** Data from D graphed to show direct comparison of SA4, Ext6, Cue-primed reinstatement and Pellet+Cue-primed reinstatement (***p<0.0001, *p<0.05). Bars indicate mean (±SEM) for each session while dots represent individual rats. AP, anterior posterior; ac, anterior commissure; NAcc, nucleus accumbens core.

### Operant behavior

*Operant chambers:* We trained and tested rats in Med-Associates (Fairfax, VT) operant chambers (#ENV-008-VPX) enclosed in sound attenuating chambers. Each operant chamber was equipped with a non-retractable lever that served as the inactive lever (#ENV-110M) to the left of the magazine (ENV-200R2M-6.0). The magazine contained a photobeam array (ENV-254-CB) to detect head entries and was connected via polyethylene tubing (Everbilt, HKP001-PVC012) to a pellet dispenser (ENV-203M-45) outside the operant chamber. To the right of the magazine was a retractable lever (ENV-112CM) with a LED stimulus light (ENV-221M) above it that served as the active lever.

*Magazine training:* We trained rats to retrieve food from a magazine in the operant chamber for 3 days for 40 trials per day (∼20 min/day). The start of training was indicated by the onset of white noise. After a variable amount of time (ITI average = 35 sec) two highly palatable food pellets (LabDiet 5TUL) were delivered into the magazine paired with a 4-sec light cue. On the last day of magazine training, we habituated the rats to the fiber optic cable. They were connected to a fiber optic patch cable (Thor labs Custom: FP-400URT, 0.5NA, 0.4m length, FT023SS tubing with a 2.5mm stainless steel ferrule) via a ceramic connector (Thor Labs, ADAF1-5) and the cable was passed through the top of the box and attached to a steel arm with a counterweight (Med Associates, PHM-110-SAI). We recorded during this last session to verify a photometry signal (see below).

*Food self-administration (SA):* We trained rats to self-administer food pellets during four daily sessions (SA1-SA4, ∼55 min/day for 4 days) with each session containing 75 trials. Like magazine training, the start of the session was indicated by the onset of white noise. Following this, the start of a trial was indicated by the extension of the active lever into the operant chamber. The lever was kept extended for 20 sec or until the rat pressed. If the rat responded on the active lever, it immediately retracted and resulted in the delivery of 2 pellets and start of a 4-sec light cue above the lever. If a rat failed to respond within 20 sec the lever was retracted and there was no consequence. The inter-trial time was variable with an average of 25 sec (minimum 20 sec, maximum 30 sec). Both active lever presses and magazine entries were recorded as our measure of food SA acquisition. Responses on the inactive lever were recorded but had no consequence. During magazine and SA training, water was provided ad libitum in the operant chamber. Photometry measures were made throughout all sessions as detailed below under *<u>Fiber photometry recordings.</u>*

*Extinction training:* Extinction training (Ext) began the day after SA4 and was identical to food SA, except that active lever presses no longer resulted in pellet delivery or cue light presentation. Six daily sessions (75 trials/session) were performed (Ext1-6).

*Reinstatement tests:* All rats received two reinstatement tests after completion of extinction training. The first test (cue-primed reinstatement; performed on the day after Ext6) began with the onset of white noise and a 4-sec cue light presentation. Subsequently, the trial began with lever extension; during the trial, active lever presses resulted in cue light illumination but no pellet delivery. On the next day, rats underwent a second reinstatement test that was identical to the first except that the initial priming event at the start of the session entailed delivery of a pellet plus presentation of the light cue (pellet+cue reinstatement). We included this to see if reward+cue prime would produce a more robust effect than priming with cue alone.

*Behavioral analysis:* Both active and inactive lever presses on the last day of SA (SA4), last day of extinction (Ext6), cue reinstatement, and pellet+cue reinstatement were compared via mixed effects analysis performed in GraphPad Prism (version 10.2.2) with an assumption of sphericity (equal variability of differences). To test for an effect of session on lever pressing, we set fixed effects of session (SA4, Ext6, Cue, Pellet+cue) and lever (active or inactive), with a random effect of subject. We made simple effect comparisons comparing active and inactive responding across sessions separately and corrected for multiple comparisons using a Holm-Šídák correction. Family-wise alpha threshold for confidence was set to 0.05. Full model output can be found in Tables S1-5. Possible sex differences were not analyzed as this study only contained 11 total subjects and therefore was not powered to detect sex effects.

### Fiber photometry recordings

*Recording parameters:* Fiber photometry was performed mainly as described in our previous study using DA sensors alone (Weber et al., 2024b). Prior to all recordings, fiber optic cables were bleached at 200 mA for 8 h overnight. On the morning of the recording day, the LED power level and DC were adjusted so that the 488 nm LED, 405 nm LED, and 560 nm LED all had a power output of 40 µW at the end of the fiber optic patch cable as measured using a Thor labs power meter (PM100D+). For recordings, we passed excitation wavelengths (488 nm, 405 nm, and 560 nm) from the TDT RZ10x system via TDT patch cables (200 µm core, 2 m length, 0.5 NA). These TDT cables then connected to a Doric Minicube (FMC6) which in turn was coupled to two Thor fiber optic patch cables. Emissions (555-570 nm,460-490 nm, and 580-680 nm) were received via the same patch cable and then decoupled at the Minicube and returned to the RZ10x via TDT response cables (600 µm, 2 m length, 0.5 NA). We acquired these emissions via the RZ10x photodetectors, digitized at 6 kHz, and recorded in the TDT Synapse Software (version 95-44132P) on WS4 at frequency 330 Hz for 465 nm, 210 Hz for 405 nm, and 450 Hz for 560 nm. All recordings had a 6 Hz lowpass filter and a 9.5 V clip threshold. During behavior sessions photometry recording was continuous for the duration of the session (all 75 trials; variable ITI of ∼52-55 min).

*Data analysis:* Fiber photometry data were analyzed using the analysis suite GuPPy created by the Lerner Lab (Northwestern University), which can be downloaded on their <u>GitHub page</u>. The details of the data manipulations used to generate the GCaMP8s and rGRAB_DA3m signal traces are described in (Sherathiya et al., 2021). In brief, for each recording session, we applied least-squares linear fit to align the same 405 nm isosbestic channel to both DA and calcium signal channels and used fluctuations in the isosbestic channel to account for florescence changes in the signal channels that are not a result of ligand binding. Any disconnects during the recording, defined by a rapid substantial shift (>10 mV) in average mV, were snipped prior to fitting the traces and any nose pokes within those snips were excluded from further analysis. Then we calculated a change in fluorescence measure (ΔF/F=Signal-Fitted Control/Fitted Control) and applied z-score normalization to control for between-session and between-rat differences in virus expression or recording. Finally, we averaged normalized traces across behavioral event replicates for each rat, and then averaged by session.

For DA and calcium transients associated with lever presses, first we extracted z-scored ΔF/F traces within a 15-sec window around each active lever press and averaged all traces to generate an average response for the rat during each test. We identified significant DA and calcium transients using continuous threshold bootstrapping methods (Jean-Richard-Dit-Bressel et al., 2020). We selected a consecutive threshold of 1000 for our 6 kHz acquisition, which equates to 0.167 s, based on recommendation from the code developer given our sampling rate. Using this analysis, we identified instances in our recorded traces where the z-score is 95% likely to not be at baseline and marked these epochs with horizontal bars above the traces. In addition to this analysis, we performed permutation tests on the bootstrapped distributions.

In tandem with this analysis, we also performed the classical hypothesis driven approach in which we selected time windows before and after our behavioral event of interest and calculated the Area Under the Curve (AUC) for the z-scored ΔF/F trace within this time window. Using these AUC values, we performed two-tailed paired t-tests comparing SA1 vs SA4, Ext01 vs Ext06, and Ext06 vs each reinstatement test. We assumed a Gaussian distribution for all t-tests and set our confidence level to 95%.

*Methodological considerations for sensor multiplexing:* Despite simultaneously recording both MSN calcium and DA, we did not make direct quantitative comparisons due to considerations detailed here. First, while we selected GCaMP8s and rGRAB_DA3m due to their reported brightness and similar kinetics, but there are small differences in kinetics which could impact our results (Zhang et al., 2023; Zhuo et al., 2023). Furthermore, measurements of these parameters are made typically in cultured neurons with controlled amounts of ligand. All our recordings are made *in vivo*, and of particular note, measure changes taking place in either in an intracellular (calcium) or extracellular (dopamine) environment. The relative abundance of calcium or DA can change the kinetics of these sensors, so we avoided comparing onset and offset of calcium and dopamine directly. Another consideration is that we normalized to baseline in all our peri-event analyses and the level of calcium or DA at our defined baseline could affect subsequent analyses. For instance, if extracellular DA is very low during baseline then any phasic release will show as a prominent peak, whereas if there is more baseline activity for calcium then any phasic increase or decrease may be harder to detect. This makes direct comparisons of the two types of transients difficult to interpret, leading us to limit our interpretations of the relationship between DA and calcium signals.

Another challenge of sensor multiplexing, particularly with green and red channels, is that many red fluorophores show emission when excited with blue light. We used a blue light channel to record an isosbestic signal for our green channel, but this light was consistently applied throughout recording and therefore no transient changes should be due to blue light excitation. However, a consideration with the isosbestic channel is signal bleed from the green GCaMP into the red GRAB_DA. We applied the same blue isosbestic recording (405 nm excitation wavelength) to both green and red channels, and it is known that, with a strong green signal, negative peaks may appear in the isosbestic recording as 405 nm is not the true isosbestic point of most GFP biosensors. Thus, when creating the ΔF/F signal, it is possible that peaks or dips are slightly amplified. When comparing isosbestic and signal channels for the same sensor this is typically not an issue but given that we applied a blue isosbestic recording to a different sensor it is possible some of the GCaMP signal may be reflected in our GRAB_DA traces. However, we believe this to be unlikely given how different these transients are in shape and in magnitude across different phases of the experiment. Furthermore, when piloting this analysis, a subset of data were processed without the inclusion of an isosbestic channel and the data appeared largely unchanged.

### Histology

Upon completion of experiments, rats were euthanized via lethal injection of Fatal Plus (Covetrus, #35946) diluted to 80 mg/kg with sterile saline. Once animals no longer displayed reflexive motor responses, they were perfused transcardially first with 1x phosphate buffer saline (PBS) followed by 4% formaldehyde/1% methanol in 1x PBS, pH ∼7, at a rate of ∼100 mL/s over 5 min. Brains were extracted and allowed to rest in the formaldehyde solution for up to 24 h. Brains were transferred to 1x PBS with 0.01% sodium azide and then sliced on a vibratome (Leica VT1000s; frequency 8, speed 70, blade DORCO plat. 5T300) at 60 µm. Slices were kept in a 24 well plate in 1x PBS with 0.01% sodium azide at 4°C until the start of immunohistochemistry.

*Immunohistochemistry:* We performed immunohistochemistry (IHC) on tissue from photometry experiments to amplify the GFP and mApple signals prior to imaging to confirm virus expression and placement. We began IHC with three 30-min washes, first in 1x PBS (diluted from 10x PBS, Quality Biological, 119-069-151) and then twice in 1x PBS with 0.5% (v/v) Triton-X100 (Electron Microscopy Sciences, #22140). After this we permeabilized the tissue for 2 h in 1x PBS with 0.5% (v/v) Triton-X100, 20% (v/v) DMSO (Sigma Aldrich, #276855), and 2% (w/v) Glycine (Sigma Aldrich, #G8898) at room temperature (RT) on a rocking shaker. Next, we blocked tissue in 1x PBS with 0.5% (v/v) Triton-X100, 10% (v/v) DMSO, and 6% (v/v) Normal Donkey Serum (NDS, Jackson Immuno Research, 017-200-121) for 2 h at RT on a rocking shaker. After blocking we incubated the tissue overnight at RT in 1:1000 Anti-GFP (Aves, GFP-1010) and 1:1000 Anti-DsRed2 (Santa Cruz Biotech, SC-101526) in 1x PBS with 0.5% (v/v) Tween-20 and 0.01% (w/v) Heparin (Sigma Aldrich, #H3393-100KU) with 3% NDS and 10% DMSO on a rotator. The following day, we washed the tissue 3 times for 30 min in 1x PBS with 0.5% (v/v) Tween-20 (Thermo Scientific, #J20605-AP) and 0.01% (w/v) Heparin before incubating in 1:250 Anti-chicken-488 (Jackson Immuno Research, 775-546-155) and Anti-mouse-647 (Jackson Immuno Research, 715-606-150) in 1x PBS with 0.5% (v/v) Tween-20 and 0.01% (w/v) Heparin with 3% NDS overnight at RT on a rotator. The third day, we performed two 30-min washes in 1x PBS with 0.5% (v/v) Tween-20 and 0.01% (w/v) Heparin before beginning a final wash in 1x PBS (30 min). Slices were kept in 1x PBS with 0.01% sodium azide at 4°C until being mounted onto Superfrost Plus slides (Fisher Scientific, Cat# 1255015) and coverslipped (Fisher Scientific, 12541026) using Vectashield Vibrance with DAPI (Vector Labs, H-1800-10).

*Imaging:* Images were acquired using a Leica DMi8 inverted microscope equipped with an ORCA-Flash4.0 LT+ Digital CMOS camera (Hamamatsu). LASX Premium Software was used for image acquisition and ImageJ was used for analysis.

## Results

Two weeks prior to the start of behavioral testing, rats received either bilateral (n = 8) or unilateral infusions (n = 3) of a viral cocktail of GCaMP8s and GRAB_rDA3m into the NAcc. In the same surgery, a fiber optic cannula was implanted above the virus injection site. Rats were allowed to recover for 1 week and then experienced 1 week of restricted chow access in the home cage. A representative image of virus co-expression of GCaMP8s and GRAB_DAr3m (showing the cannula track) is provided in Fig. 1B, whereas cannula placements are summarized in Fig. 1C.

### Behavioral results

We trained rats to lever press for delivery of 2 highly palatable pellets paired with a 4-sec light cue. Rats readily acquired this behavior, with the majority reaching 100% trial completion (75 trials) after 4 days of SA training (Fig. 1D; this panel shows responding during SA training and also during extinction training and reinstatement tests). We then put rats through extinction training in which active lever presses no longer resulted in a food reward or a light cue. After 6 sessions of extinction rats had suppressed their responding on the active lever compared to the last day of self-administration (Fig. 1E; *t*_59_ = 14.77, *p*< 0.0001). They also reduced responding on the inactive lever, a permanently extended lever associated with no consequence throughout all training and testing (Fig. 1E; *t*_59_ = 3.779, *p* = 0.0022). Once lever pressing had been extinguished, we tested the rats for cue-primed reinstatement. During this ∼55 min test, lever presses delivered the same 4-sec light cue previously paired with food reward. The session began with a single non-contingent presentation of the light. Rats increased responding for the cue compared to the last day of extinction (Fig. 1E; *t*_59_ = 2.980, *p* = 0.0125) with no effect on inactive lever responding. On the next day, we performed a second reinstatement test identical to the first except that the initial light cue also was paired with non-contingent delivery of a pellet, acting as a reward+cue prime. As in the previous reinstatement test, rats also exhibited increased responding on active but not inactive levers (Fig. 1E; *t*_59_ = 2.572, *p* = 0.0282). See Table S1 for the full statistical output.

### Photometry overview and analysis of food self-administration training sessions

We simultaneously recorded both MSN calcium transients (Fig. 2) and DA release (Fig. 3) in the NAcc on all days of the regimen. For all traces, we examined a 15-sec window around the active lever press. GCaMP and DA traces are plotted to compare the first and last day of food SA (Fig. 2A and 3A), the first and last day of extinction (Fig. 2B and 3B), the last day of extinction and the cue-primed reinstatement (Fig. 2C and 3C), and the last day of extinction and the pellet+cue-primed reinstatement (Fig. 2D and 3D). Above all the traces are horizontal lines of matching color that indicate periods where the 95% confidence interval does not cross baseline (i.e., z-score = 0) for the consecutive threshold period determined with a continuous threshold bootstrapping method (Jean-Richard-Dit-Bressel et al., 2020; Liu et al., 2020; Yau and McNally, 2022). Time periods during which permutation testing revealed differences between sessions are indicated by a light gray line.

**Figure 2.**
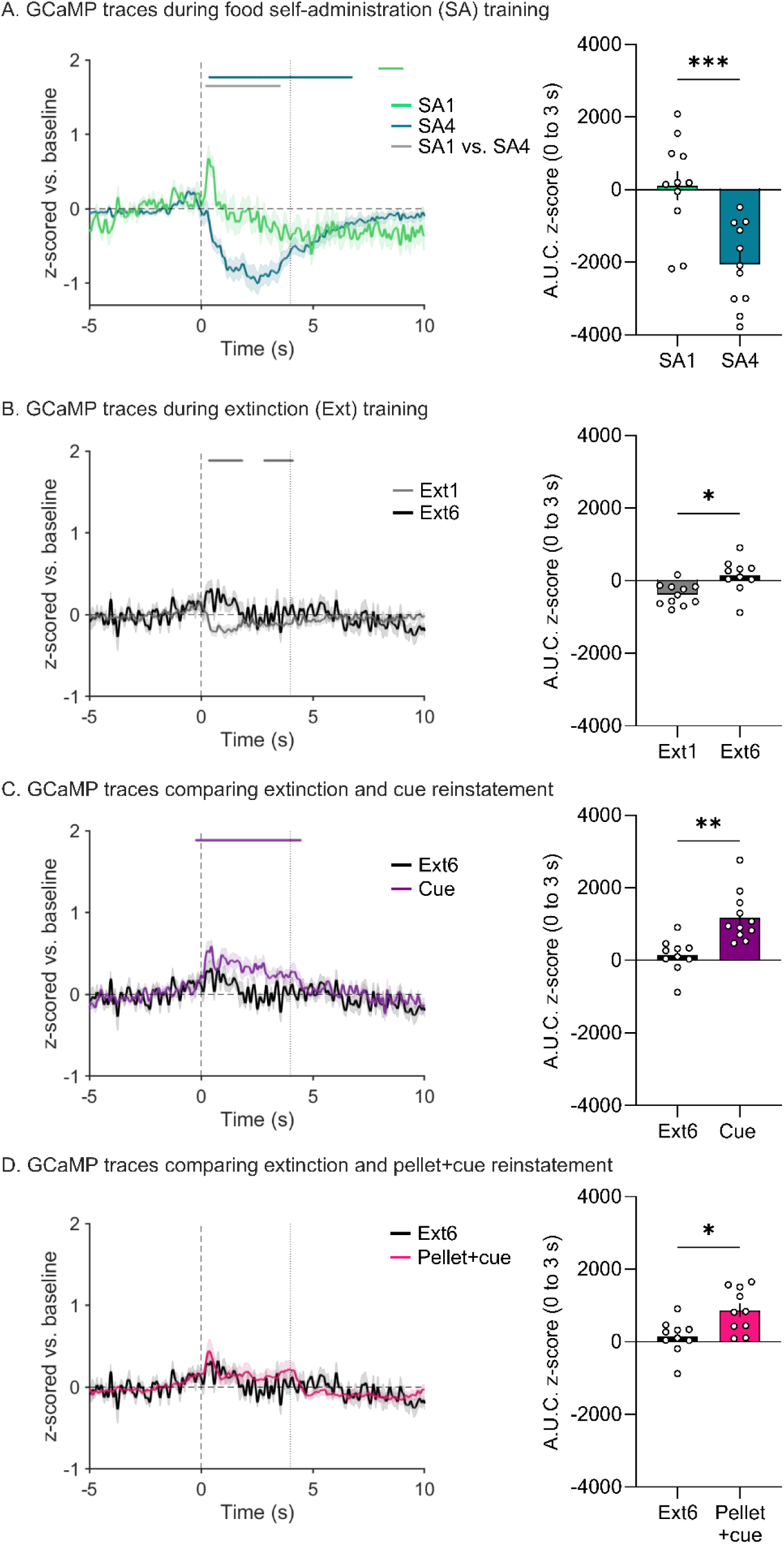
Fiber photometry recordings of NAcc calcium signals across food self-administration, extinction, and reinstatement sessions. **A.** *Left:* z-scored mean GCaMP traces time-locked to active lever presses in first session of self-administration (SA1) and the last session of self-administration (SA4) normalized to a baseline period (-5 to 0 s). SEM is shown in shaded area around the mean. The black, vertical dashed line at time 0 s indicates the lever press with the gray, vertical dashed line at time 4 s indicating the end of the light cue. The matching-colored lines above the traces show periods in the 15-s window during which bootstrapping indicates 95% confidence that the mean is not equal to zero (baseline level). *Right:* Area under the curve for the traces shown in *Left*. Bars show mean (±SEM) for 3 s after the lever press while dots indicate individual rats. **B-D.** These panels show GCaMP traces, as described in **A,** during the first session of extinction (Ext1) and the last session of extinction (Ext6) (**B**), the last session of extinction (Ext6) and cue-primed reinstatement (Cue) (**C**), and the last session of extinction (Ext6) and pellet+cue-primed reinstatement (Pellet+cue) (**D)**.

**Figure 3.**
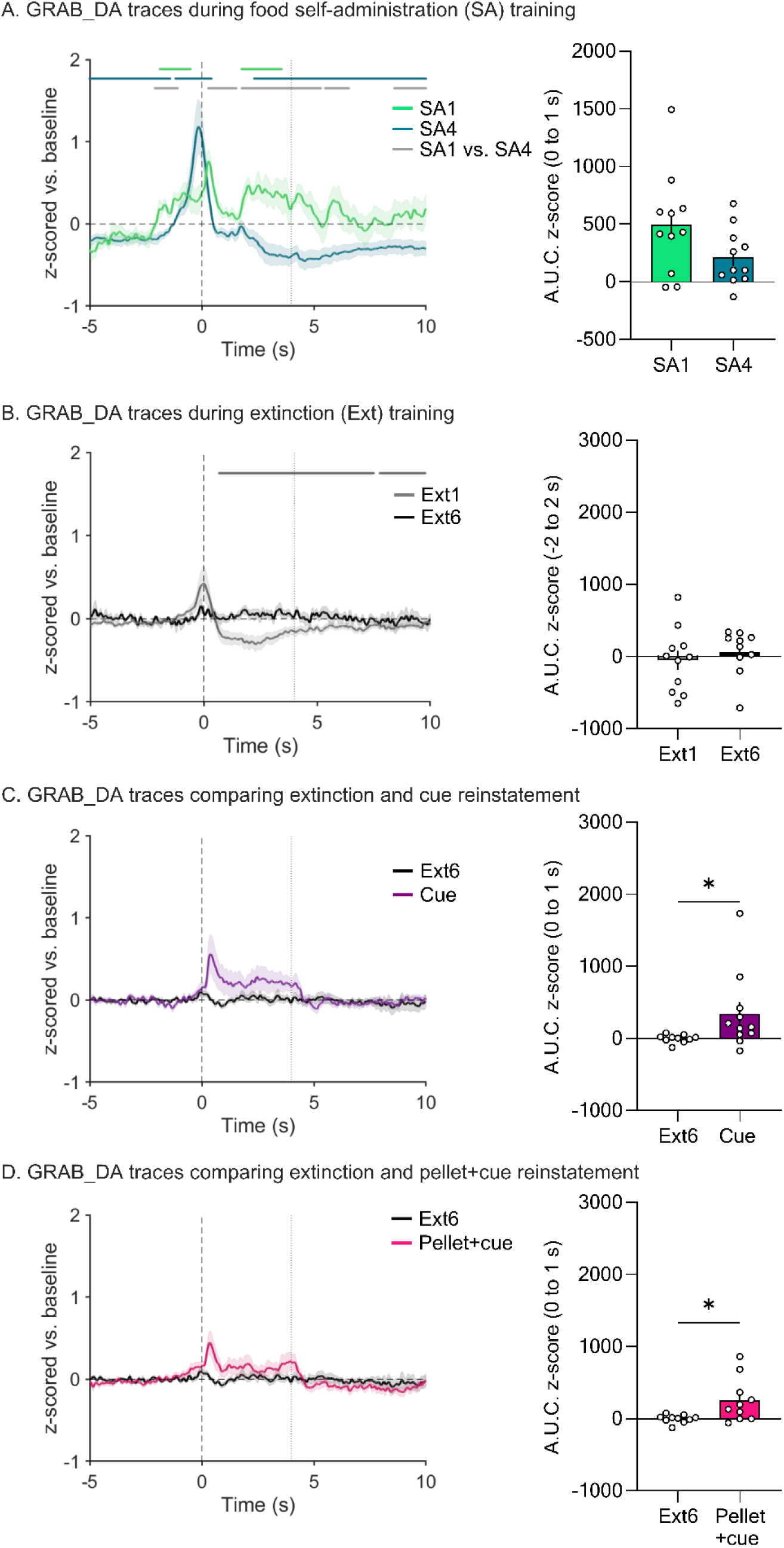
Fiber photometry recordings of DA transients in the NAcc across self-administration, extinction, and reinstatement sessions. **A.** *Left:* z-scored mean DA traces time-locked to active lever presses in first session (SA1) and last session (SA4) of self-administration normalized to a baseline period (-5 to 0 s). SEM is shown in shaded area around the mean. Black, vertical dashed line at time 0 s indicates the lever press. Gray, vertical dashed line at time 4 s indicates the end of the light cue. The matching-colored lines above the traces identify significant transients, i.e., periods in the 15-s window (-5 to 10 s) during which bootstrapping indicates 95% confidence that the mean is not equal to zero (baseline level). *Right:* Area under the curve for the traces shown in *Left*. Bars show mean (±SEM) for time window 2 s prior and 2 s after the lever press while dots indicate individual rats. **B-D.** These panels show GRAB_DA traces, as described in **A,** during the first session of extinction (Ext1) and the last session of extinction (Ext6) (**B**), the last session of extinction (Ext6) and cue-primed reinstatement (Cue) (**C**), and the last session of extinction (Ext6) and pellet+cue-primed reinstatement (Pellet+cue) (**D)**. *p<0.05.

Focusing on comparison of food SA sessions 1 and 4 (SA1 and SA4), analysis of GCaMP traces with bootstrapping showed the development of a significant suppression of MSN calcium levels as training progressed (Fig. 2A left; statistical analysis in Table S2). We quantified the area under the curve (AUC) for the time period 0 to 3-s and compared SA1 to SA4 with a paired t-test to confirm this decrease (Fig. 2A right; *t*_10_=5.143, *p*=0.0004; full model output in Table S3). Extinction and reinstatement data are detailed in separate sections below. However, we note that AUC data for calcium signals on each SA day, as well as Ext1-6 and reinstatement tests, are presented together in Fig. S1 for ease of comparison (0 to 3-s window). This was not done for DA data because different windows were required to capture temporally shifting DA responses across the experiment, as described below.

A different pattern emerged for DA traces (Fig. 3; full model output in Tables S4 and S5). Upon lever press, we observed a small increase in DA in SA1 that appeared to become more robust and shift earlier in time by SA4 (Fig. 3A). During SA4, this DA peak was followed by a persistent dip below baseline (Fig. 3A left). These differences between SA1 and SA4 were corroborated by bootstrapping analysis (Fig. 3A left). In addition, we analyzed AUC for the 0 to 1-s window to attempt to capture the reduction in DA signal following the lever press as it shifted earlier in time (i.e., out of this window; Fig. 3A right) and then analyzed peak height in the-5 to 0-s window to capture the gradual emergence from SA1 to SA4 of a DA signal preceding the lever press (Fig. S2). In both cases, we note that animal-to-animal variability made it difficult to select an ideal window to capture each change, and thus the reduction in Fig. 3A only trended (p=0.09) although the emergence in Fig. S2 was statistically significant.

Overall, fiber photometry results obtained during SA training suggest that MSN activity, as measured by calcium, decreases from basal levels upon a lever press that results in food reward and that this develops over the course of training. In contrast, DA release associated with active responding rises from baseline levels and this peak becomes more robust and moves to precede the lever press. A potential explanation is that DA release begins to track lever entry (which precedes the lever press) as the rats learn to perform the task.

### Photometry analysis of extinction sessions

We performed the same analysis described above for the first and last sessions (Ext1 and Ext6) of extinction training. For GCaMP, bootstrapping analysis indicated a small post-press decrease during Ext1 that is eliminated by Ext6 (Fig. 2B). When comparing the AUC for these two sessions with a paired t-test, this change is significant (Fig. 2B right, Table S3; *t*_9_=3.000, *p*=0.0150). In fact, there was very little GCaMP response at any point in Ext6. Of note, one rat made no responses during Ext6 and therefore could not be included in analysis. For DA, bootstrapping analysis showed a small positive peak preceding the lever press on Ext1, similar to what was observed during SA4 (and thus perhaps reflecting anticipation of reward), but this was not evident during Ext6 (Fig. 3B left). Analysis of AUC during the window of the positive peak (-2 to 2 s, 0 being the lever press) did not detect any peak in either Ext1 or Ext6 with no significant difference between the two sessions (Fig. 3B right). Bootstrapping also indicated a post-press dip below baseline during Ext1 which dissipated by Ext6 (Fig. 3B left). Together, these findings indicate a loss of DA responding over the course of extinction training. Furthermore, these data suggest that the calcium and DA responses observed in early extinction are not due to the motor action of the lever press (as the motor action is equivalent across extinction sessions) but rather due to the reward and learned operant response outcome. However, when comparing these two sessions it is important to keep in mind that the number of trials completed in Ext6 is greatly reduced compared to Ext1, which could potentially have affected our analysis.

### Photometry analysis during reinstatement tests

Reinstatement tests allowed us to further examine NAcc MSN calcium and DA responses in a reward seeking context. When examining MSN calcium via GCaMP, bootstrapping analysis indicated a small increase from baseline that overlapped with the light cue presentation in the cue-primed (Fig. 2C left) but not pellet+cue-primed tests (Fig. 2D left). When assessed with AUC, however, this increase in calcium was significant for both reinstatement tests compared to the last extinction session (paired t-tests; cue: *t*_9_ = 3.413, *p* = 0.0077; pellet+cue: *t*_8_ = 2.393, *p* = 0.0436) (Fig. 2C right and 2D right). For DA transients, there was an apparent increase for cue-primed reinstatement and pellet-primed reinstatement (positive peak overlapping with cue presentation) (Fig. 3C left and Fig. 3D left). While neither was considered a likely peak by bootstrapping analysis (Fig. 3C left and 3D left), analysis with AUC indicated a significant increase in magnitude of the DA response during both reinstatement tests compared to the last extinction session (*t_9_*=2.589, *p*=0.0293 and *t_8_*=2.519*, p*=0.0359 for cue-and pellet+cue-primed reinstatement, respectively) (Fig. 3C right and 3D right).

## Discussion

Here we used sensor multiplexing to assess the relationship between NAcc MSN activity (using calcium as a proxy) and DA transmission during food self-administration, extinction, and reinstatement. Our finding of a large dip in the z-scored GCaMP signal during food self-administration agrees with prior observations of decreased NAc MSN neuronal firing rate during food consumption (see below). However, this dip develops over the course of training and is also observed in the first extinction session, although it disappears over subsequent sessions. These observations suggest a relationship to task learning and not solely food consumption. In the reinstatement phase, an elevation of MSN calcium parallels increased cue-induced responding. In the same rats, we observe a DA peak that initially follows lever pressing but moves earlier in time, and appears to increase in size, as self-administration training progresses. This peak is reduced in the first extinction session and eventually disappears with extinction learning, although a DA response is restored during reinstatement. Overall, these findings show a complex relationship between MSN calcium and DA transmission during food self-administration training and subsequent extinction, although increases in both calcium and DA transmission occur during reinstatement. We note that technical considerations preclude direct comparison of the magnitude and timing of calcium and DA signals (see Materials & Methods).

### Role of NAcc MSN and DA transmission in food consumption

The role of NAcc MSN activity in food intake has been investigated using *in vivo* electrophysiology. This previous work found that a subset of neurons increase their firing rate during operant responding for food but exhibit a pause in firing during consumption that is necessary for consummatory behavior (Carelli et al., 2000; Nicola et al., 2004; Roitman et al., 2005; Frazier and Mrejeru, 2010; Krause et al., 2010). Although we acknowledge that calcium is only a proxy for neuronal activity, our calcium data seem to parallel this prior work, with a negative transient occurring after the lever press. This could be partially due to reward consumption, but there is also likely a learning component. Thus, if the observed calcium decrease was simply due to reward consumption it likely would be unchanged across SA sessions, but instead we find that this decrease in calcium becomes more pronounced with training. This could reflect higher trial completion, and therefore a higher number of rewards eaten, but as we only analyzed completed trials, we believe this to be unlikely. Supporting a role for learning, a study observing a pause during consumption found that it was modulated depending on the presentation of a predictive cue (Nicola et al., 2004). Likely, the lever extension at the start of the trial acts as our predictive cue or ‘DS’ and could be modulating the decrease in calcium signaling as the task is learned. Supporting this, a small decrease from baseline upon lever press remains evident in Ext1. In this session there is zero consumption, indicating that some portion of this decrease is due to other aspects of behavior. Future studies manipulating NAcc function will be required to isolate the behavioral component encoded by this decrease from basal calcium. For example, it would be interesting to determine the effect of exciting these neurons during the extinction phase.

DA has long been thought to be essential for reward learning, but the exact nature of its role is debated. The classic view of DA is that it encodes discrepancies between the prediction of reward as a result of behavior and the actual consequences, termed “reward-prediction error” (Keiflin and Janak, 2015). In this model of DA as a learning signal, it would be expected that DA responses would be elevated early in food self-administration training (when the rat is learning the relationship between a lever press and reward delivery), decrease later in training (as the reward becomes more predicted), and increase in extinction (when the response-outcome contingency changes again). That pattern of DA responding is not reflected in our data. This could be due to our analysis of session averages, and perhaps if we investigated within SA1 and Ext1 we would find evidence of classical RPE signaling. A more recent theory of DA’s significance proposes that it acts as a value signal and, particularly in the NAc, high DA indicates that effortful work will be beneficial (Berke, 2018). This tracks better with our observations, with DA increasing as the rat completes more trials and becomes more assured of the value of the lever press, and decreasing in Ext1 when the value of that work reduces. Other theories propose DA as a signal of perceived saliency (Kutlu et al., 2021) or a tool for retrospective learning (Jeong et al., 2022), both of which would also track with our observations of DA transients during food SA and extinction.

### Role of NAcc MSN and DA transmission in food seeking

The reinstatement phase isolates the value of the reward-associated cue and its ability to invigorate behavior. We found significantly increased calcium transients in both the cue-primed and pellet+cue-primed reinstatement tests compared to the last session of extinction (AUC analysis for both tests and bootstrapping for the former). Likewise, there was an increased DA response during both reinstatement tests compared to the last session of extinction (AUC analysis). Thus, calcium and DA exhibit opposite responses during SA (above) but track together during reinstatement tests. Some studies design their task to investigate both the cue in isolation and the reward response by having the cue precede the operant action. These studies found increased activity of NAc neurons during the cue period which was facilitated by DA (Yun et al., 2004; du Hoffmann and Nicola, 2014). Furthermore, intra-NAc DA receptor antagonism with SCH23390 or raclopride attenuated cue-induced food reinstatement (Guy et al., 2011). These results, along with ours, suggest that the ability of cues to promote food seeking depends upon MSN activation (presumably driven by glutamate) enhanced by neuromodulatory effects of DA.

### Role of D1 and D2 MSN in observed responses

Our recordings sampled all neurons in the NAcc, but it is important to consider the cell types that are contributing to this general population signal. The principal neurons of the NAcc are GABAergic MSNs that express either D1 or D2 DA receptors (D1 MSN, D2 MSN). D1 MSN have often been implicated in driving motivated behaviors whereas D2 MSN are less involved or oppose such behaviors, although current theories emphasize the importance of balance and interactions between these pathways (Burke et al., 2017; Bariselli et al., 2019; Allichon et al., 2021). We expect that both D1 and D2 MSN are represented in our population signal, but may contribute differently to observed signals. However, even for our most robust effect (a decrease in the calcium signal from baseline after lever pressing that intensifies from SA1 to SA4; Fig. 2A), it is difficult to speculate as to the relative involvement of D1 vs D2 MSN. In agreement with our findings, many early studies reported that a subset of MSN show inhibition during initiation and maintenance of sucrose consumption (Nicola et al., 2004; Roitman et al., 2005; Taha and Fields, 2005, 2006; Krause et al., 2010). Importantly, electrical stimulation of NAc regions containing inhibited neurons interrupts consumption, indicating that their inhibition plays a permissive role (Krause et al., 2010). Among subsequent studies that distinguished D1 and D2 MSN and measured their activity (using electrophysiology or fiber photometry) during palatable solution consumption, one study found that lateral habenula-projecting shell D1 MSN reduce activity to permit consumption while D2 MSN do not reliably change activity(O’Connor et al., 2015), while others observed D2 MSN inhibition associated with consumption (Guillaumin et al., 2023) or observed inhibition of both MSN subtypes after a lick but differences preceding the lick (Walle et al., 2024). The different results likely reflect substantially different paradigms and methodologies and, together with other analyses performed as part of these studies, suggest that a complex interplay of D1 and D2 MSN governs consummatory behavior. In the future, it would be interesting to repeat our experimental paradigm with GCaMP selectively expressed in D1 vs D2 MSN. However, the main point of the present study was to demonstrate the feasibility of a multiplexing approach to assess MSN activity simultaneously with dopamine levels.

### Conclusions

NAcc neuronal activity as measured by intracellular calcium was progressively reduced during food self-administration training, while the inverse was observed for DA signaling. Both responses were reduced on the first day of extinction before disappearing (no detectable transient) by the last day of extinction. When measured during cue-primed and pellet+cue-primed reinstatement, significant increases in calcium and DA were observed relative to the last day of extinction. In addition to providing new information about calcium and DA dynamics in this food reward paradigm, this study also served to validate sensor multiplexing as a technique that can be applied in the future to studies of natural rewards and drugs of abuse.

## Acknowledgements

This work was supported by F31 DA057063 (SJW) and OHSU startup funds (MEW). We thank Dr. Rajtarun Madangopal for assistance with MedPC code, Dr. Venus Sherathiya for assistance with GuPPy, and Randall Olson for assistance with MATLAB code.

## Supplementary Information (Figures S1 and S2, Table S1)

**Figure S1.**
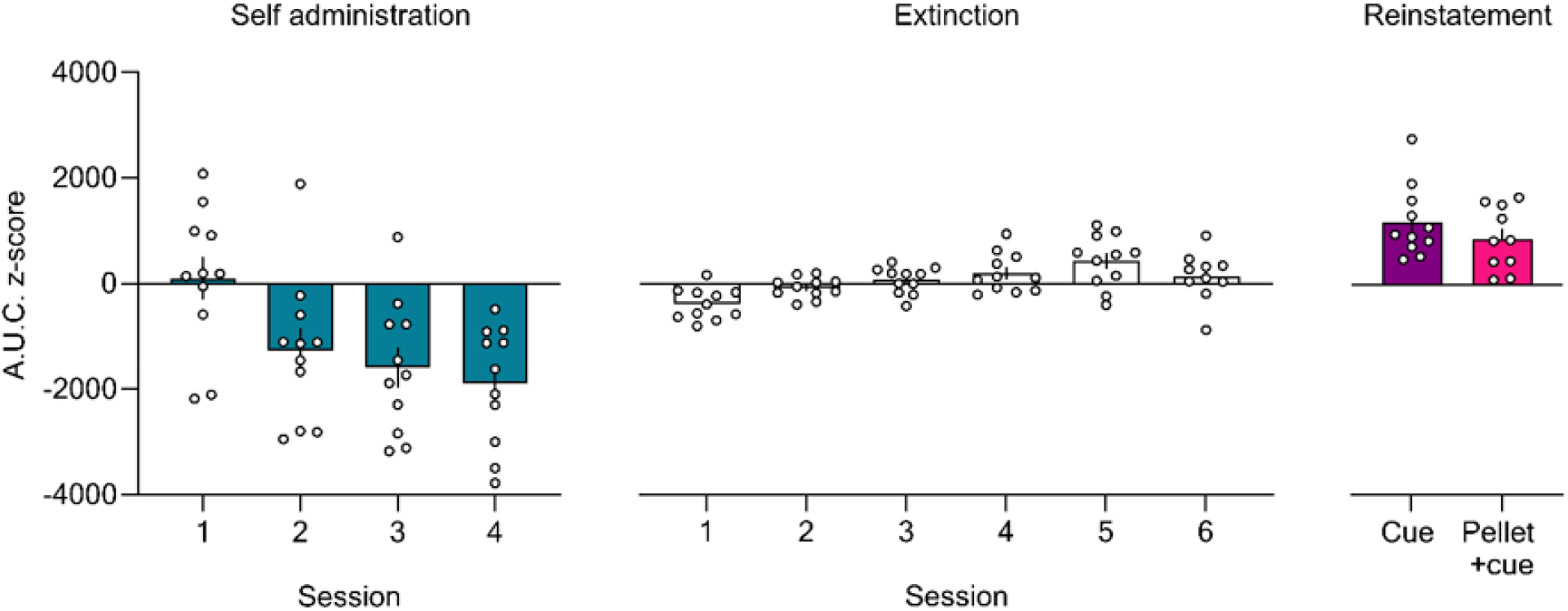
Area under the curve (AUC) for GCaMP traces across all recorded sessions. The time period analyzed corresponds to that shown in Fig. 2, i.e., 0 to 3 s after the lever press. Bars show mean (±SEM) AUC while dots indicate individual rats.

**Figure S2.**
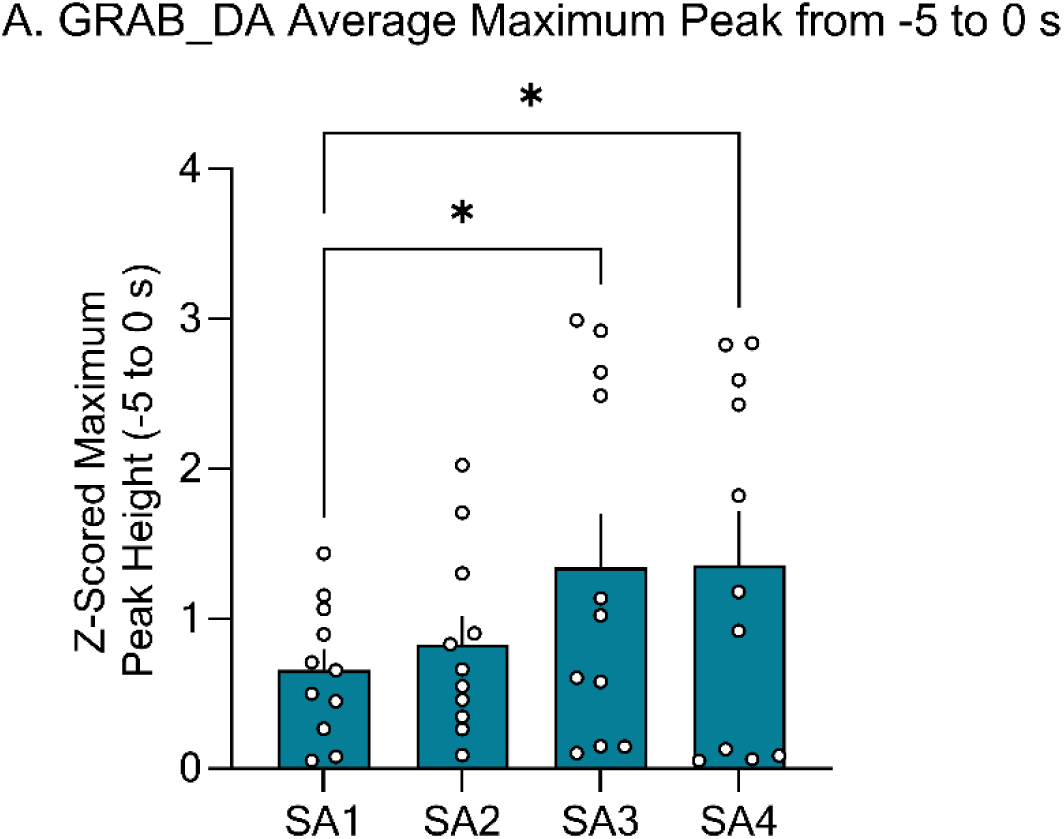
Z-scored maximum peak height for GRAB_DA traces across all SA sessions. The time period analyzed corresponds to the 5 sec prior to the lever press. Data were analyzed with a one-way ANOVA. Bars show mean (±SEM) Maximum Peak while dots indicate individual rats.

---

**Table S1.** Statistical output for behavioral data in Figure 1.

| Expt phase | Measure | Fixed effects | F-value | P-value | Significant? | Figure |
| --- | --- | --- | --- | --- | --- | --- |
| SA4, Ext6, Cue, Pellet+cue | Lever presses (n = 11, 6M/5F) | <u>Session x Lever</u> |  |  |  | 1E |
| | | Session | $F_{3,30}=68.67$ | <0.0001 | **** | |
| | | Lever | $F_{1,10}=88.86$ | <0.0001 | **** | |
| | | Session x Lever | $F_{3,29}=23.85$ | <0.0001 | **** | |
|  |  | <u>Holm-Šídák<sup>†</sup></u> |  |  |  |  |
|  |  | Inactive |  |  |  |  |
| | | SA4 vs. Ext6 | $t_{59}=3.779$ | 0.0022 | ** | |
| | | SA4 vs. Cue | $t_{59}=2.995$ | 0.0159 | * | |
| | | SA4 vs. Pellet+cue | $t_{59}=3.542$ | 0.0039 | ** | |
| | | Ext6 vs. Cue | $t_{59}=0.6811$ | 0.8739 | n.s. | |
| | | Ext6 vs. Pellet+cue | $t_{59}=0.2376$ | 0.8805 | n.s. | |
| | | Cue vs. Pellet+cue | $t_{59}=0.4500$ | 0.8805 | n.s. | |
|  |  | Active |  |  |  |  |
| | | SA4 vs. Ext6 | $t_{59}=14.77$ | <0.0001 | **** | |
| | | SA4 vs. Cue | $t_{59}=11.79$ | <0.0001 | **** | |
| | | SA4 vs. Pellet+cue | $t_{59}=12.24$ | <0.0001 | **** | |
| | | Ext6 vs. Cue | $t_{59}=2.980$ | 0.0125 | * | |
| | | Ext6 vs. Pellet+cue | $t_{59}=2.527$ | 0.0282 | * | |
| | | Cue vs. Pellet+cue | $t_{59}=0.4535$ | 0.6518 | n.s. | |

<sup>†</sup>adjusted P-values reported for post hoc comparisons. M, male, F, female.

**Table S2.** Statistical output for bootstrapping analyses in Figure 2.

| Expt phase | Measure | Factors in analysis | Time 95% CI $\neq 0$ | Significantly different? | Figure |
| --- | --- | --- | --- | --- | --- |
| SA | GCaMP response to lever press, z-scored trace (n=11) | <u>Bootstrapping</u> |  | 0.235 to 3.51 s | 2A, left |
|  |  | SA1 | n.s. |  |  |
|  |  | SA4 | 0.375 to 6.69 s |  |  |
| Extinction | GCaMP response to lever press, z-scored trace (n=11) | <u>Bootstrapping</u> |  | n.s. | 2B, left |
|  |  | Ext1 | 0.394 to 1.84 s |  |  |
|  |  | Ext6 | n.s. |  |  |
| Extinction/Reinstatement | GCaMP response to lever press, z-scored trace (n=11) | <u>Bootstrapping</u> |  | n.s. | 2C, left |
|  |  | Ext6 | n.s. |  |  |
|  |  | Cue test | -0.214 to 4.46 s |  |  |
| Extinction/Reinstatement | GCaMP response to lever press, z-scored trace (n=11) | <u>Bootstrapping</u> |  | n.s. | 2D, left |
|  |  | Ext6 | n.s. |  |  |
|  |  | Pellet+cue test | n.s. |  |  |

**Table S3.** Statistical output for AUC GCaMP fiber photometry data in Figure 2.

| Expt phase | Measure | Comparison | T-value | P-value | Significant? | Figure |
| --- | --- | --- | --- | --- | --- | --- |
| SA | AUC (n = 11) | SA1 vs. SA4 | $t_{10}=5.143$ | 0.0004 | *** | 2A, right |
| Extinction | AUC (n = 10) | Ext1 vs. Ext6 | $t_9=3.000$ | 0.0150 | * | 2B, right |
| Extinction/<br>reinstatement | AUC (n = 10) | Ext6 vs. Cue | $t_9=3.413$ | 0.0077 | ** | 2C, right |
| Extinction/<br>reinstatement | AUC (n = 9) | Ext6 vs. Pellet+cue | $t_8=2.393$ | 0.0436 | * | 2D, right |

**Table S4.** Statistical output for bootstrapping analyses in Figure 3.

| Expt phase | Measure | Factors in analysis | Time 95% CI $\neq 0$ | Significantly different? | Figure |
| --- | --- | --- | --- | --- | --- |
| SA | GRAB_DA response to lever press, z-scored trace (n=11) | <u>Bootstrapping</u> |  |  |  |
|  |  | SA1 | -1.82 to -0.507 s, 1.79 to 3.55 s | -2.01 to -1.07 s, 0.285 to 1.53 s, 1.79 to 5.23 s, 5.56 to 6.83 s, 8.55 to 10.0 s | 3A, left |
|  |  | SA4 | -5.00 to -1.41 s, -1.13 to 1.77 s, 2.38 to 10.0 s |  |  |
| Extinction | GRAB_DA response to lever press, z-scored trace (n=11) | <u>Bootstrapping</u> |  |  |  |
|  |  | Ext1 | to -2.89 s, 0.71 to 7.44 s, 7.77 to 9.75 s | n.s. | 3B, left |
|  |  | Ext6 | n.s. |  |  |
| Extinction/<br>Reinstatement | GRAB_DA response to lever press, z-scored trace (n=11) | <u>Bootstrapping</u> |  |  |  |
|  |  | Ext6 | n.s. | n.s. | 3C, left |
|  |  | Cue test | n.s. |  |  |
| Extinction/<br>Reinstatement | GRAB_DA response to lever press, z-scored trace (n=11) | <u>Bootstrapping</u> |  |  |  |
|  |  | Ext6 | n.s. | n.s. | 3D, left |
|  |  | Pellet+cue test | n.s. |  |  |

**Table S5.** Statistical output for AUC GRAB_DA fiber photometry data in Figure 3.

| Expt phase | Measure | Comparison | T-value | P-value | Significant? |  | Figure |
| --- | --- | --- | --- | --- | --- | --- | --- |
| SA | AUC (n = 11) | SA1 vs. SA4 | $t_{10}=1.970$ | 0.0856 | n.s. | | 3A, right |
| Extinction | AUC (n = 10) | Ext1 vs. Ext6 | $t_9=0.4708$ | 0.6490 | n.s. | | 3B, right |
| Extinction/<br>reinstatement | AUC (n = 10) | Ext6 vs. Cue | $t_9=2.589$ | 0.0293 | * | | 3C, right |
| Extinction/<br>reinstatement | AUC (n = 10) | Ext6 vs. Pellet+cue | $t_9=2.519$ | 0.0359 | * | | 3D, right |

**Table S6.** Statistical output for Peak Height GRAB_DA fiber photometry data in Figure S2.

| Expt phase | Measure | Fixed effects | F-value | P-value | Significant? | Figure |
| --- | --- | --- | --- | --- | --- | --- |
| SA | Peak Height (n = 11) | Session | $F_{3,30}=4.396$ | 0.0112 | * | S2A |
|  |  | Holm-Šidák† |  |  |  |  |
| | | SA1 vs. SA2 | $t_{30}=0.7093$ | 0.4836 | n.s. | |
| | | SA1 vs. SA3 | $t_{30}=2.843$ | 0.0205 | * | |
| | | SA1 vs. SA4 | $t_{30}=2.903$ | 0.0205 | * | |

